# A fluid droplet harvests the force generated by shrinking microtubules in living cells

**DOI:** 10.1101/2024.09.09.612121

**Authors:** Katherine Morelli, Sandro M. Meier, Angela Zhao, Madhurima Choudhury, M Willis, Yves Barral, Jackie Vogel

## Abstract

The energy-consuming dynamic instability of microtubules generates significant forces which are thought to be harnessed to move large cargos in cells. However, identification of mechanisms which can capture the force released during microtubule depolymerization to move large loads has been elusive. In this work we show that a biomolecular condensate provides an elegant solution to this problem. Using live cell super-resolution microscopy, we directly observe that budding yeast +TIP bodies are nanoscale droplets with classic fluid-like behaviors which accumulate type V myosin (Myo2) at their surfaces. We find that conserved self-oligomerization interfaces in the protein Kar9 tune the biophysical properties of the viscoelastic +TIP body and its ability to efficiently move the mitotic spindle. Our findings introduce a paradigm for how forces generated by microtubule dynamics are harnessed in cells and open a frontier of research on nanoscale biomolecular condensates in their native environment.

## Introduction

Microtubules are essential for the movement of many cargos in cells. They have a well-known role as dedicated tracks for the processive walking of motor proteins, e.g. dyneins and kinesins that are loaded with cargos. In eukaryotes, this mechanism is responsible for moving large complex cargos such as vesicles, mitochondria, and mRNPs^1^. A second and less well-understood mechanism by which microtubules move cargos is through their end-on attachments, such as during the critical movement of chromosomes during mitosis^2^.

End-on attachments are thought to capture and harness the significant forces generated by microtubule dynamic instability^2^. The microtubule polymer assembles in an energy consuming process that stores mechanical energy. GTP-bound tubulin dimers assemble end to end into protofilaments. These protofilaments assemble through lateral contacts to form hollow tubes. Since GDP-bound protofilaments prefer to bend outward, the hydrolysis of GTP allows strain to build up in the shaft as the microtubule grows. During microtubule shrinkage, protofilaments are thought to peel outward in individual power strokes. Altogether, they can generate force in the 10-30 pN range^3–5^. This force, if captured, equals that of ensembles of many motor proteins^6^. However, how cells harvest these forces to move larger cargos is largely unknown^7,8^.

The most well-characterized mechanism for capturing the force released by microtubules is the end-on attachment of the DASH/Dam1 ring complex of the budding yeast kinetochore. In the proposed model, the microtubule-encircling DASH/Dam1 ring complex simultaneously captures the power strokes of all outwardly peeling protofilaments from a single microtubule to move the kinetochore^5^, but no analogous ring has been identified in metazoans^9^, suggesting that other types of mechanisms must exist. The question of how large cargos can remain attached to dynamic microtubule +ends, particularly since any protein-protein interaction to specific tubulin dimers will need to be reformed as the dimers are lost during shrinkage, remains largely unresolved.

Interactions between microtubule plus ends and the cell cortex are critical for positioning the mitotic spindle in all eukaryotic cells, and is executed by deeply conserved +TIP proteins and molecular motors. Mitotic spindle positioning in budding yeast provides a unique opportunity to resolve how cells can use forces generated by a small number of shrinking microtubules to move large cargos such as the spindle. During mitosis, the spindle is moved into position along the mother-bud axis by a few astral microtubules^10^. In budding yeast, all astral microtubules are nucleated from the spindle pole bodies (centrosomes), and all dynamics happen at their plus-ends, which probe the cytoplasm^11^. Metaphase spindle positioning requires the microtubule plus-end tracking protein (+TIP) Kar9, which couples microtubules to the polarized actin network emanating from the bud tip ^10,12,13^. At anaphase onset, one spindle pole body must enter the bud to ensure proper inheritance of chromosomes. This can occur via Kar9-dependent cortical anchoring during microtubule shrinkage, which pulls the spindle into place^10,13^. In the absence of Kar9, a second Kar9-independent pathway which positions the spindle via dynein becomes essential^14^. The molecular arrangement of Kar9-dependent cortical attachments and how they can transduce force has not been resolved.

The components of the Kar9 pathway were recently found to form a highly cohesive structure called the +TIP body which tracks shrinking microtubules. It contains Bim1 (EB1 homolog), Bik1 (CLIP-170 ortholog), and Kar9, which form a highly multivalent and network of redundant, homo- and heterotypic interactions among themselves (Figure 1A). In vitro and indirect evidence in vivo suggest that the +TIP body is a phase-separated condensate (Figure 1A)^15^. The +TIP body is an excellent candidate for harvesting the force generated by microtubule shrinkage^15^. Phase-separated condensates can perform work through capillary phenomena like wetting^16^, which may explain how the +TIP body tracks shrinking microtubules. Capillary phenomena or viscoelasticity can store force through resistance to deformation, which may explain how cortical attachments transduce force^17^. However, measuring these properties for nanoscale condensates in vivo is challenging because of their dynamic nature and size close to the resolution limit of traditional light microscopy.

**Figure 1.**
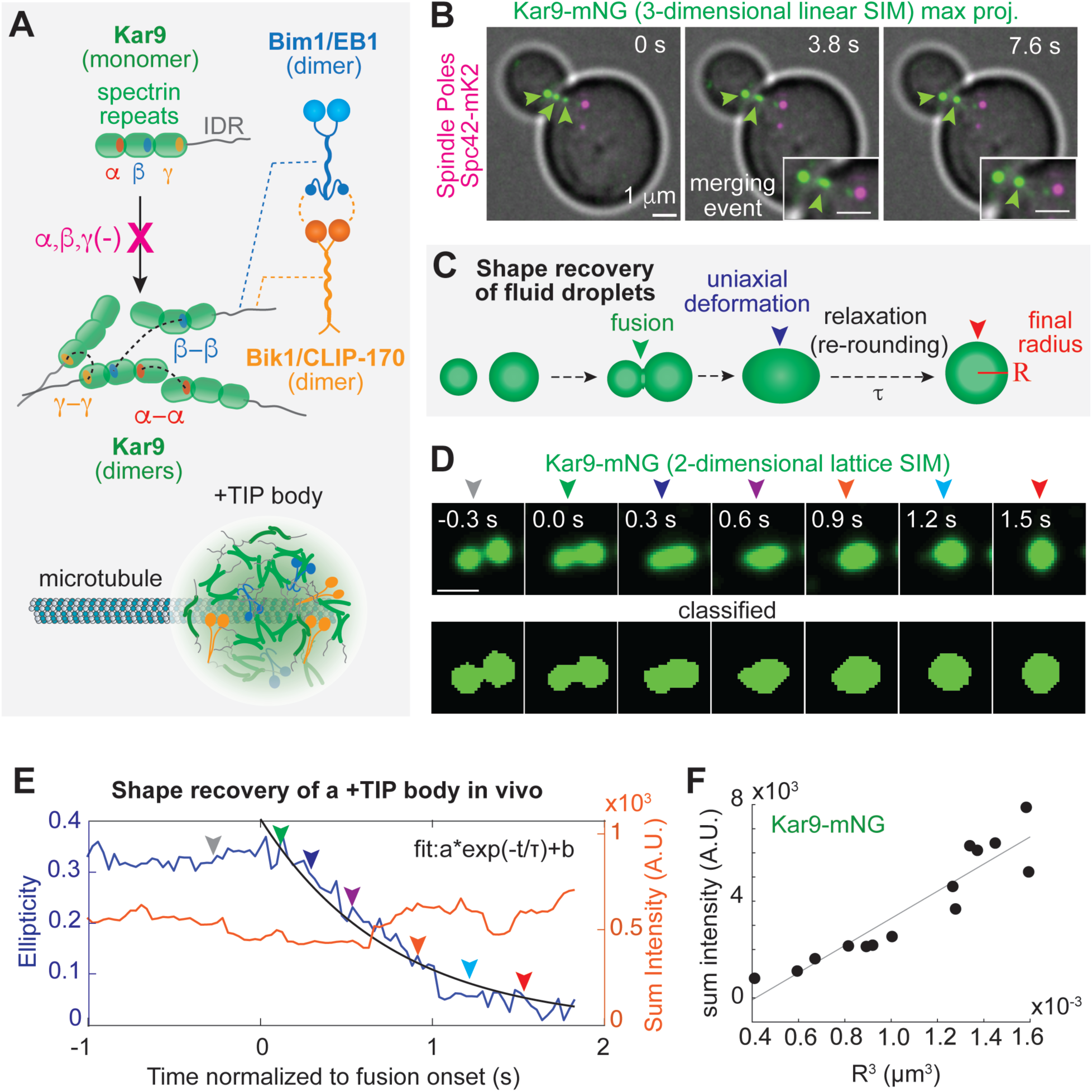
+TIP body shape recovery dynamics in vivo. (A) Schematic of the multivalent Kar9-Bim1/EB1-Bik1/CLIP-170 network and interaction with astral microtubules. Known protein-protein interactions depicted as dotted lines. (B) 3D maximum intensity projection of live-cell 3D linear SIM time series (timestep = 3.8 seconds) imaging of Kar9-mNG and spindle poles labeled with Spc42-mK2, overlaid with BF image of the same field of view, which displays a merging event. Scale bar is 1 µm. (C) Theory of shape recovery of fluid droplets upon fusion (green arrow) and uniaxial deformation (blue arrow). The characteristic relaxation time (ι−) measures the timescale of re-rounding. The characteristic length scale is R, the radius of the final droplet. If internal η is much larger than external, then 1/R ≈ η/γ, where γ is surface tension. This is the inverse capillary velocity. (D) Above: Live-cell 2D lattice SIM time series (31 frames per second) imaging of Kar9-mNG which displays a merging event. Arrows indicate positions in trajectory from E). Below: Pixel classification used for shape analysis. Scale bar is 0.5 µm. (E) Plot of ellipticity and sum fluorescent intensity (A.U.) over time from data in D). Arrows indicate corresponding timepoints between (D) and (E). Ellipticity is defined as (L-W)/(L+W) where L = length and W = width. (F) Sum fluorescent intensity (A.U.) vs R^3^ (R = final radius) for +TIP bodies resulting from merging events.

Because of these di]iculties, making direct links between properties of nanoscale condensates and their function in vivo has been recognized as an important frontier in the field of biomolecular condensates^16^.

To examine the force-harvesting capabilities of the +TIP body in living cells, we need direct measurements of its biophysical properties and function in vivo^16,18^. Here, we report direct measurements of the fluid behavior of the yeast +TIP body using high-speed super-resolution microscopy and investigate its role in cargo movement. Our findings that demonstrate the +TIP body is a force-transducing nanoscale fluid droplet and that these measurements can be made in living cells.

## Results

### Direct measurements of Kar9 fluid-like behavior in living cells

We have observed budding yeast +TIP bodies undergoing fusion and fission events consistent with the behavior of fluid droplets^15^ (Figure 1B). The classical behavior of droplets from physics has been applied to the study of sub-cellular biology since pioneering work identified the liquid behavior of P granules^19^ and nucleoli^20^. We reasoned that high-speed lattice structured illumination microscopy (lattice SIM) could visualize the shape of dynamic, nanoscale +TIP bodies during merging events. This approach would allow us to directly measure whether merging events follow the expected behavior of fluid droplets (Figure 1C).

We acquired 2-dimensional lattice SIM time series in living cells of +TIP bodies labeled endogenously with Kar9 fused C-terminally to mNeonGreen (mNG,as in^21^, from^22^), achieving a lateral resolution of ∼100 nm and effective frame rate of 31 frames per second. From these acquisitions, we identified events in which initially separate foci merged into a single focus (Figure 1D). The Kar9-mNG fluorescent signal was segmented using pixel classification (Figure 1D). Events with relatively constant total fluorescent intensity (Figure 1E, Figure S1B), indicating that they remained in focus during the event, were used for analysis. To quantify the shape recovery of +TIP bodies upon a merging event, we measured the ellipticity of the segmented +TIP body through the merging event until the shape re-rounds (defined as ellipticity < 0.1, Figure 1E, Figure S1A, C, see Methods). In most events the +TIP body re-rounds within five seconds (Figure 1E, Figure S1A, C; n=13/14) upon merging. This indicates that Kar9-mNG molecules rapidly reorganize into a larger sphere as expected for a fluid. This outcome distinguishes +TIP bodies from solid-like objects which would stick to each other without re-rounding.

Our pixel segmentation threshold was chosen to capture approaching +TIP bodies as one continuous shape before they appear as a single punctum (Figure 1D). We reasoned that using our lattice SIM 2-dimensional time series, the exact moment at which the surfaces of the two +TIP bodies come into contact cannot be directly observed. However we can infer this moment by looking for a changepoint in the shape trajectory. Remarkably, such a changepoint is visible in our shape trajectories (Figure 1E, Figure S1A, C). If +TIP bodies are viscous droplets in cells, we expected re-rounding into larger spheres should evolve over time as an exponential decay function, which is driven by surface tension and opposed by viscosity^23^. Indeed, we found that the terminal behavior of the ellipticity trajectories after the changepoint is well approximated by an exponential decay function (Figure 1E, Figure S1A). Additionally, for the +TIP bodies resulting from fusion, total intensity scales with the cube of the radius of the final shape, as expected for 3-dimensonal droplets (Figure 1F). Altogether these results indicate that at the 0.1-1s timescale, +TIP bodies behave as fluid droplets in living cells.

### +TIP bodies can be much larger than microtubule +ends in living cells

The size of the +TIP body has functional implications; we speculate that larger diameter +TIP bodies could accelerate cortical capture, host multiple microtubules, and accumulate surfactants. If the +TIP network proteins coat microtubule plus-ends fully (diameter 25nm), we expect that +TIP bodies cannot be smaller than around 40 nm.

However, the existence of fusion events in addition to the radii detected during these events suggests that some +TIP bodies may be much larger (Figure 1).

We reasoned that the moment of fusion onset, when droplets accelerate toward each other, indicates when droplets first interact and can be used to provide an alternative measure of length scale (Figure 2A). We reason that the distance between the two peaks of fluorescent intensity from the merging +TIP bodies, at the moment of fusion onset, is an estimate of sum of their radii R_1_ + R_2_ (Figure 2A). We measured the distance between the centers at the moment of fusion by fitting a 2-mean gaussian function (Figure 2B). The peak distance at the moment of fusion onset measured this way yields similar measures to the final diameter, which ranged from 150-250 nm (Figure 2C). Our measurements likely report the upper bound of the size distribution, since they are measured from +TIP bodies which undergo observable fusion events. We also measured an event with two consecutive fusions (Figure 2D). Both the peak distance and final diameter increased together as expected for the two consecutive fusions (Figure 2E). Together, our measurements indicate that the +TIP body can become as large as 8-fold greater than the diameter of a single microtubule (Figure 2F).

**Figure 2.**
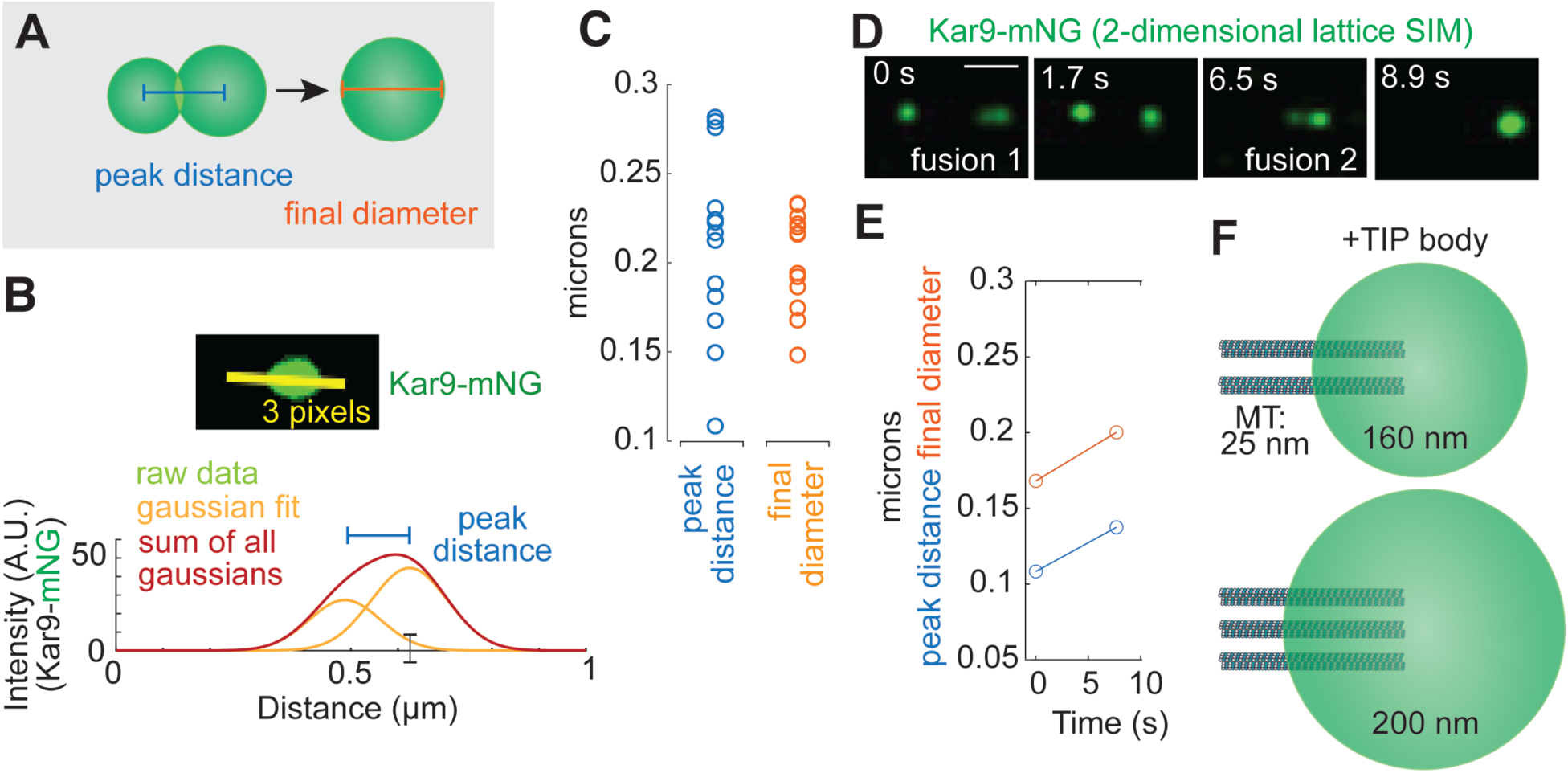
Shape recovery dynamics measure +TIP body size. (A) Schematic of two measures of length scale. For two spheres, the sum of their radii R_1_ + R_2_ (peak distance) is related to final diameter D of the sphere which is the sum of their volumes by the relation 1.26*(R_1_ + R_2_) ≤ D ≤ 2*(R_1_ + R_2_). (B) Above: Live-cell 2D lattice SIM time series (31 frames per second) imaging of Kar9-mNG which displays fusing +TIP bodies at the moment of fusion onset. Overlay displays 1µm, 3 pixel width linescan used for analysis. Below: Plot of raw data, gaussian fits, and sum of all gaussians used to to measure the peak distance across the linescan. (C) Plot comparing +TIP body length scale measured by final diameter and peak distance for wild type cells. (D) Live-cell 2D lattice SIM time series (31 frames per second) imaging of Kar9-mNG which displays two consecutive fusing events. Scale bar 0.5 µm. (E) Plot comparing final diameter and peak distance following each fusion event in D, time normalized to the onset of the initial event. (F) Schematic of potential +TIP body dimensions and microtubule distribution in vivo based on C,D, and E.

### Myo2 motors accumulate at the +TIP body surface

The plus ends of astral microtubules position the spindle at the bud neck through their interaction with actin cables that originate from the bud cortex. This interaction requires Kar9, which is both a cargo of the budding yeast actin-base motor Myo2 and a component of the +TIP body ^24^. Our results indicate that the +TIP body is a droplet which can be more than 200 nm in diameter, and thus could interact with many Myo2 motors. Since Kar9 is known to directly bind the Myo2 cargo domain^25^, we reasoned that in cells Myo2 may localize to either the surface or the dense phase of the +TIP body.

To begin addressing this question, we first reconstituted +TIP bodies by combining purified Kar9, Bim1, and Bik1 protein and applied cell lysate containing Myo2-GFP (in^26^, from^27^) tagged at the endogenous locus. Interestingly, when we induced phase separation (Figure 3A), we observed that Myo2-GFP was excluded from the bulk of the reconstituted droplets, instead appearing as puncta localized at the surface (Figure 3B). In contrast, application of cell lysate containing only GFP displayed a uniform distribution throughout reconstituted droplets (Figure 3B). This indicates that Myo2 is excluded from the bulk and instead accumulates at the surface of reconstituted +TIP droplets.

**Figure 3.**
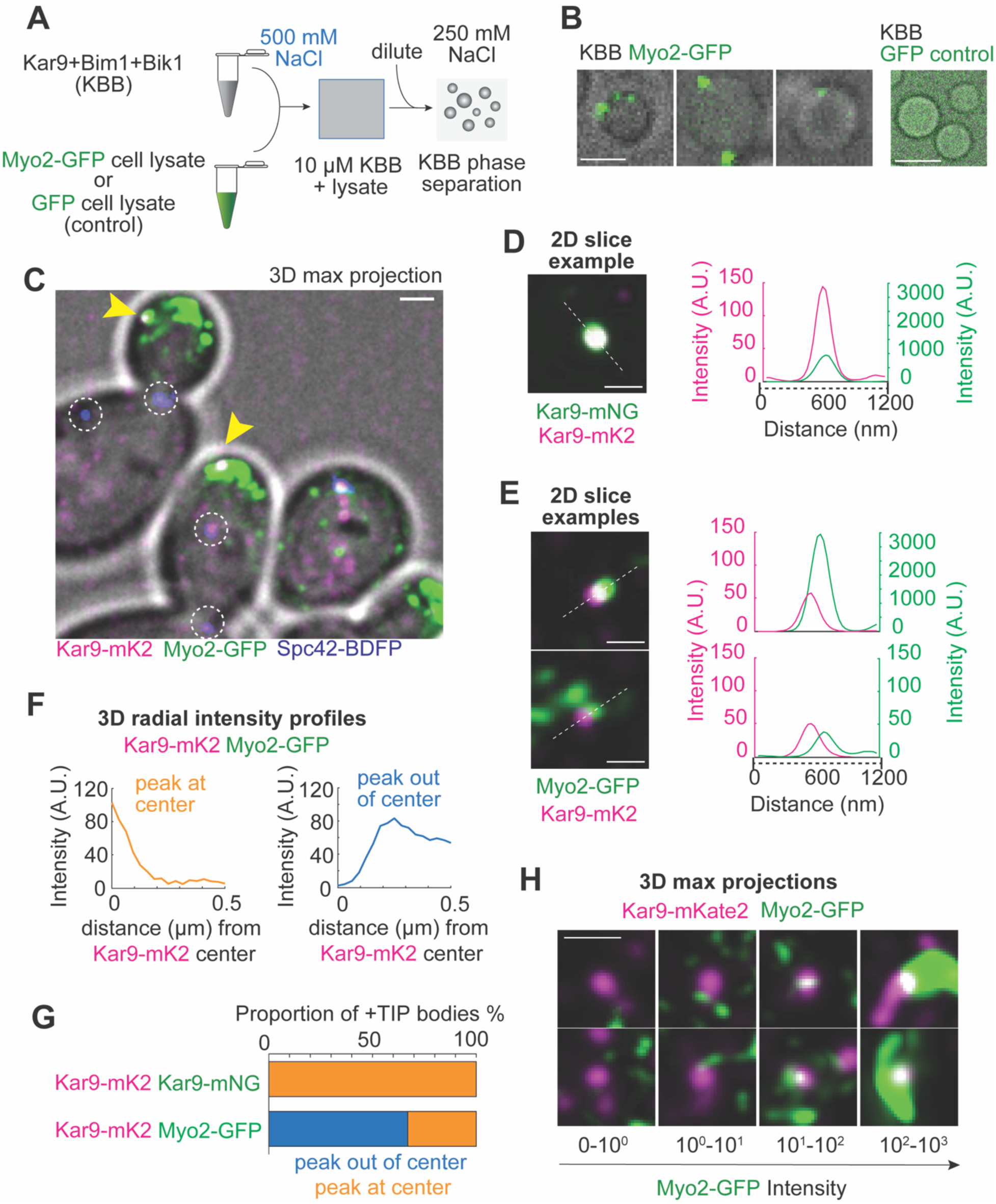
Myo2 association with +TIP bodies. (A) Schematic of in vitro droplet partitioning assay. Myo-GFP or GFP control cell lysate added to purified Kar9 + Bim1 + Bik1. (B) Confocal microscopy images from in vitro droplet partitioning assay described in (A). Scale bar 2 µm. (C) 3D maximum intensity projection of live-cell 3D linear SIM imaging of Kar9-mK2, Myo2-GFP, and spindle poles labeled with Spc42-BDFP, overlaid with BF of the same field of view. Scale bar is 1 micron. Scale bar 2 µm. Yellow arrows indicate cortical Kar9 puncta colocalized with Myo2. White dashed circles indicate spindle pole body positions. Far-red BDFP channel is manually masked to remove Myo2 signal bleed-through and to visualize spindle pole bodies more easily. (D) Live-cell 3D linear SIM imaging of Kar9-mK2, Kar9-mNG diploid cell. 2D slice taken from centroid of Kar9-mK2 signal. Scale bar 0.5 µm. Intensity profiles along a 1.2 µm 1-pixel width linescan in red and green channel from central slice are depicted. E) Live-cell imaging as in (C). 2D slice taken from centroid of Kar9-mKate2 signal. Green channel is Myo2-GFP. Scale bar 0.5 µm. Intensity profiles along a 1.2 µm 1-pixel width linescan in red and green channel from central slice are depicted. F) Plots of 3D radial normalized intensity (A.U.) in the green channel as distance from the +TIP body center increases for individual +TIP bodies. Examples depict both profile types: peak out of center, peak at center. G) Plot of proportions of +TIP bodies with peak out of center and peak at center profile types. (Kar9mK2; Kar9mNG n=9, Kar9-mK2; Myo2-GFP n = 69) H) Maximum intensity projections of 1µm depth centered around the centroid of Kar9-mKate2 signal. Images are grouped by the maximum value from radial intensity profile of Myo-GFP. Scale bar 0.5 µm.

Next, we asked whether the surface partitioning that occurs in vitro is characteristic of Myo2 interactions with +TIP bodies in living cells. To address this question, we acquired 3-dimensional linear SIM super-resolution acquisitions of +TIP bodies and Myo2 labeled endogenously using Kar9-mKate2 (mK2, from^28^) and Myo2-GFP (Figure 3C). If Myo2 binding to Kar9 is unrestricted we would expect that Kar9-mK2 and Myo2-GFP fluorescent signals would fully overlap in 2-dimensional slices of +TIP bodies. This expected overlap is shown for +TIP bodies in control diploid cells expressing both Kar9-mK2 and Kar9-mNG (Figure 3D). In contrast, we find that for many +TIP bodies, the Kar9-mK2 and Myo2-GFP signals do not fully overlap (Figure 3E).

To quantify the potential exclusion of Myo2 from the bulk of +TIP bodies, we measured the normalized radial intensity of the Myo2-GFP signal from the center of the Kar9-mK2 signal (Figure 3F). Remarkably, for the majority of +TIP bodies, the brightest point of normalized signal was not at the center, as it is for all control +TIP bodies which are dual-labeled with Kar9-mK2 and Kar9-mNG (Figure 3F-G). This suggests that Myo2-GFP is excluded from the bulk phase of +TIP bodies in living cells. This observation is consistent with the existence of a bona fide surface of +TIP bodies in living cells and their fluid-like properties. We also noted heterogeneity in the amount of Myo2-GFP colocalized with Kar9, with the peak intensity of the green channel signal of Myo2 varying by at least four orders of magnitude (Figure 3H). This indicates that +TIP bodies can host hundreds of associated Myo2 motors. Together these results suggest that +TIP bodies interact with cortical actin filaments via Myo2 motors which do not stoichiometrically bind to Kar9 but rather adhere to those on the +TIP body surface.

### Conserved oligomerization interfaces tune the biophysical properties of +TIP bodies

Our results suggest that +TIP bodies are nanoscale fluid droplets that can associate with Myo2 motors via their surfaces. This architecture suggests that the emergent biophysical properties of the +TIP body as a biomolecular condensate could play an important role in its function as a force-coupling device between dynamic microtubule ends and the actin cortex. Since fusion dynamics are sensitive to underlying material properties, we explored using them to characterize the biophysical properties of +TIP bodies.

We first quantified the timescale of fusion events by extracting the characteristic time (τ) from the exponential fit (Figure 4C). The timescale of fusion events scales with the length scale of the +TIP body radius (R), and the resulting value of τ /R is proportional to the inverse capillary velocity^19^. Reanalyzing data from Meier et al. 2023^15^ (Figure S2), we find that the inverse capillary velocity of in vitro reconstituted droplets composed of purified Bim1, Bik1, and Kar9 is strikingly similar to our in vivo measurements (Figure 5B). This indicates that the capillary velocity (∼0.5µm/s) is faster than the velocity of microtubule dynamics (∼1µm/min)^29^, consistent with a model in which capillary action contributes to +TIP bodies’ ability to track growing shrinking microtubules.

**Figure 4.**
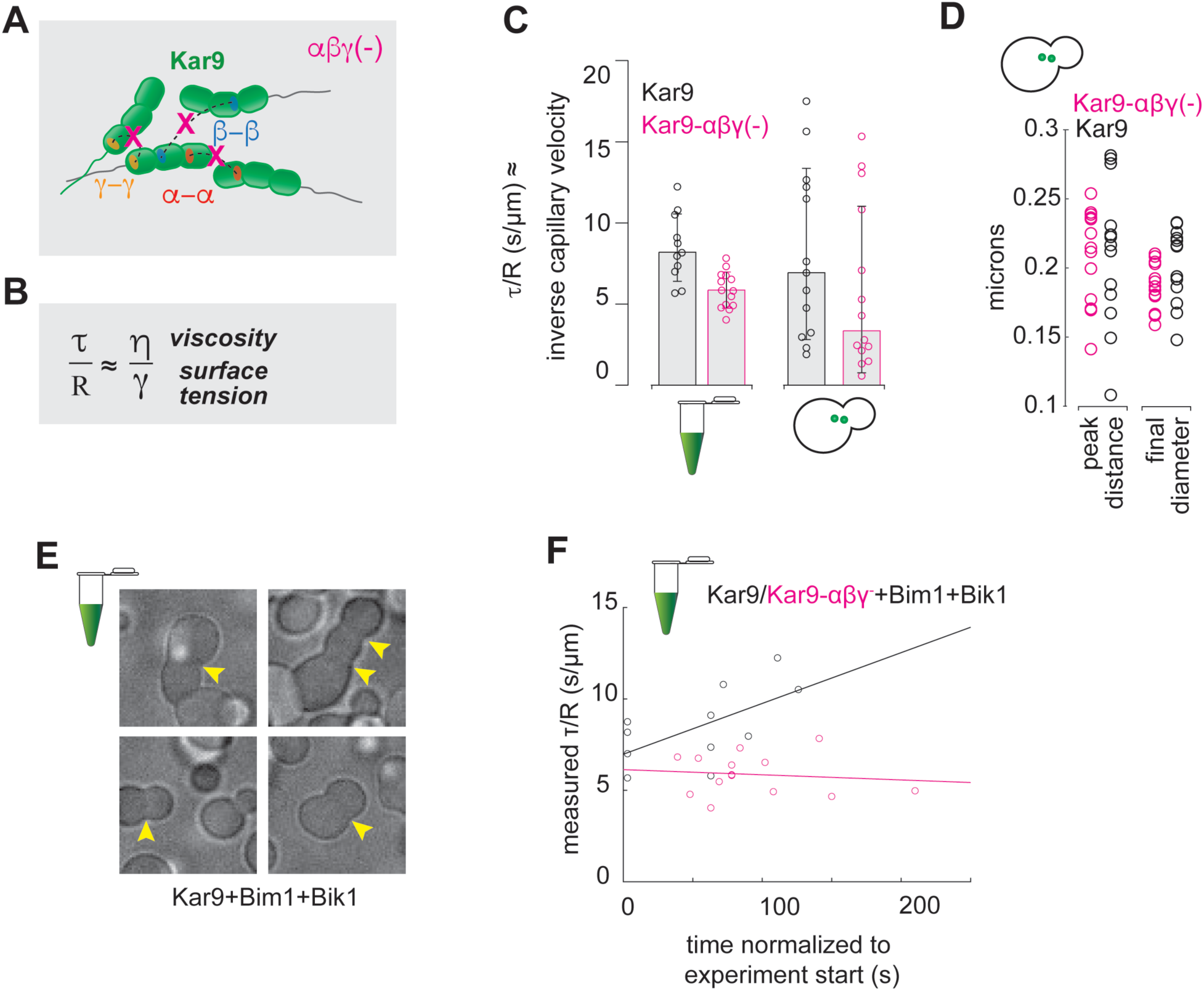
Biophysical characterization of Kar9 oligomerization interface mutant +TIP bodies. (A) Schematic of Kar9-αβγ(-) mutants. (B) The characteristic relaxation time (1) measures the timescale of re-rounding. The characteristic length scale is R, the radius of the final droplet. If internal viscosity η is much larger than external viscosity, then 1/R ≈ η/γ, where γ is surface tension. This is approximately the inverse capillary velocity. (C) Plot comparing independent τ/R (s/µm) measurements from reconstituted +TIP bodies (Kar9-HIS or Kar9-αβγ-HIS, Bim1, Bik1-HIS; data reanalyzed from^15^) and native +TIP bodies in vivo. Kar9/Bim1/Bik1 n=11 median=8.2 std=2.1, Kar9-αβγ/Bim1/Bik1 n=14, median=5.8, sd=1.1, WT n=13 median=6.9, sd= 5.3, Kar9-αβγ-: n=14 mean=3.5, sd=5.1. (D) Comparison of Kar9-αβγ(-) +TIP body size measured by final diameter and peak distance. (E) KBB droplets with stalled fusion. (F) Plot of τ/R (s/µm) measurements from reconstituted +TIP bodies (Kar9-HIS or Kar9-αβγ-HIS, Bim1, Bik1-HIS; data reanalyzed from^15^) over experimental time.

**Figure 5.**
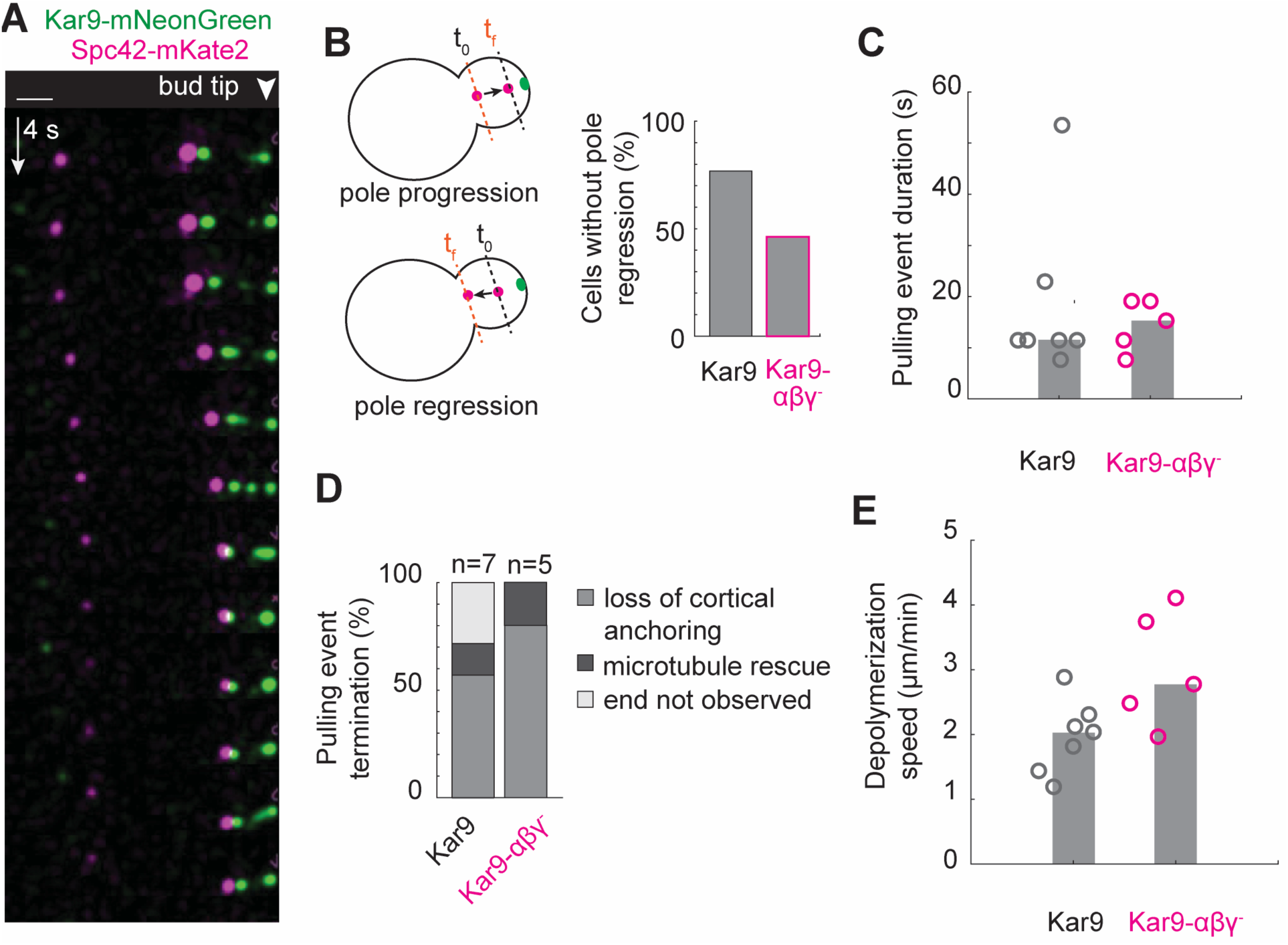
Spindle movements during depolymerization of microtubules with cortically anchored +TIP bodies. (A) 3D maximum intensity projections of live-cell 3D linear SIM time series (timestep = 3.8 seconds) imaging of Kar9-mNG and spindle poles labeled with Spc42-mK2, which displays a pulling event. Right side of image is the position of the bud tip, indicated by a white arrow. Scale bar is 1 µm. (B) Schematic of spindle pole regression measurement and plot of pole regression for wild type and Kar9-αβγ(-) cells. (C) Plot of duration of pulling events for wild type and Kar9-αβγ(-) cells. (D) Plot of pulling event termination causes for wild type and Kar9-αβγ(-) cells. F) Plot of inferred microtubule depolymerization velocity during pulling events for wild type and Kar9-αβγ(-) cells.

The structured N-terminus of Kar9 contains three self-interaction interfaces (α, β, and γ) which lie in conserved domains^15,26^ (Figure 1A). Since these interfaces are not mutually exclusive, together they can create oligomerization. We speculated that mutations in the α, β, and γ interfaces that reduce oligomerization may alter the material properties of +TIP bodies sufficiently to be detected by our methods in living cells. We observed that Kar9-αβγ(-) mutant +TIP bodies fuse in living cells. Like wild type, most resulting bodies re-round (Figure 5C). For both in vitro and in vivo mutant +TIP bodies, the measured inverse capillary velocity is lower (Figure 4A). This both affirms the reliability of our nanoscale fusion dynamic analysis in vivo and indicates that the α, β, and γ interfaces contribute to material properties in living cells.

The range these values fall in is consistent with other highly viscous biomolecular condensates with comparatively low surface tension which allows them to be stable and persistent at the mesoscopic length scales relevant in living cells ^30^. These values are also consistent with the high degree of valency among +TIP proteins within +TIP bodies^15^.

Importantly, for mutant +TIP bodies resulting from fusion events, we find that the total intensity scales with the cube of the radius (Figure S3). However, this relationship is less steep than in wild type cells, indicating that αβγ(-) +TIP bodies have less density of Kar9 in vivo. Consistent with this, using the center at fusion analysis to estimate size, we find that mutant +TIP bodies are similar in size despite having less overall intensity (Figure 5D). This reduction of Kar9 density in the αβγ(-) mutant may underlie the observed change in material properties in vivo by reducing viscosity and/or increasing surface tension. Previous work noted that +TIP bodies in Kar9-αβγ(-); bik1Δ cells undergo apparent fusion and fission events more frequently than in wild type^15^.

Interestingly, we also noted that for our in vitro reconstitution assays, we measured that the τ /R increased over time in the wild type movie but not the mutant (Figure 4E). The slowing of fusion towards the end of the wild type movie is suggestive of the competition between surface energy and elastic modulus, or in other words, viscoelasticity. This may suggest that the interaction network promoted by Kar9 oligomerization may contribute to viscoelasticity of droplets. Increase in capillary velocity over the course of minutes in these in vitro reconstitutions indicates a significant capacity for condensate aging. Since in cells we observed a minority of events with incomplete or stalled fusion (Figure 4C), we speculate that +TIP bodies maintain their fluidity and prevent condensate aging through active processes in living cells.

Taken together, we find that mutations in the α, β, and γ interfaces that reduce Kar9 oligomerization do not significantly reduce the size of +TIP bodies but rather produce +TIP bodies with altered properties in their native environment.

### Cortically anchored +TIP bodies on shrinking microtubules position the spindle

Previous work identified that mutations disrupting Kar9 oligomerization (Figure 5A) caused an increase in errors of the critical process of spindle positioning at anaphase onset^15^. We wondered whether these defects could result from a reduced ability to form force-transducing links coupling shrinking astral microtubules to the cortex during fast anaphase.

To address this question, we acquired 3-dimensional linear SIM acquisitions with Kar9-mNG and the spindle pole body component Spc42 tagged C-terminally with mKate2 (as in ^21^) with a timestep of 4 seconds. We measured the instantaneous length and growth rate of spindles and used this measure along with the movement of the spindle across the bud neck in some cases to identify cells in fast anaphase (Figure S5)^31^. We observed that cells in fast anaphase had apparent +TIP body cortical anchoring and pulling events (Figure 5A). We defined cortical anchoring as when a cortical focus of Kar9 at the bud cortex in the bud moved less than 200 nm over a window >8 seconds while the spindle pole moved towards it, indicating that the spindle pole body moved towards the cortical location as the microtubule was depolymerizing (Figure 5A). To measure the efficiency of spindle pole movements into the bud, we classified each cell by whether the spindle pole regressed or was able to maintain or advance its position relative to the tip of the bud during the 68 second observation window (Figure 5B). Only around half of Kar9-αβγ(-) cells maintained or advanced the position of the old spindle pole body during the acquisition (Figure 5B), indicating that spindle positioning is less efficient in this mutant. We next asked whether the Kar9-αβγ(-) mutant resulted in shorter pulling events. We found that pulling events in wild type cells lasted longer on average than in mutant cells (Figure 5B: Wild type mean=

18.6 s, n=7; Kar9-αβγ(-) mutant mean = 14.5 s, n=5). This indicates that wild type cells in fast anaphase spend more time in a state where the spindle is productively cortically anchored. We also noted there were two events in wild type cells whose end was not observed (Figure 5C), suggesting that an observation window of 68 seconds is limiting, and the difference may even be larger. We also noted that in the Kar9-αβγ(-) mutant cells, more events ended because cortical anchoring was lost (Figure 5D). Interestingly, we found that in Kar9-αβγ(-) mutant, microtubule depolymerization velocities during pulling events were faster (Figure 5E). Altogether, in Kar9-αβγ(-) mutant cells, cortical anchoring is less stable and depolymerization pulling events are shorter, reducing the efficiency of spindle positioning during the critical step of fast anaphase.

## Discussion

Biomolecular condensates are ubiquitous in critical eukaryotic cellular processes^32^. Extensive in silico and in vitro characterization of their remarkable emergent properties^16,33^ suggests these properties may answer many long-standing mysteries in cells. However, direct characterization of endogenous material properties, which evolved for functions in living cells, is limited. Nanoscale condensates, such as the recently discovered class of eukaryotic +TIP bodies, present additional challenges for in vivo studies because of their dynamicity and size. In this work we establish a method for measuring the properties and function of budding yeast +TIP bodies in living cells and provide insight to a long-standing question of how microtubule dynamics are harnessed in cells to do work.

Our high-resolution time series of +TIP body fusion in cells demonstrates that +TIP bodies are fluids, and their properties can be measured in cells. We restrict imaging to a 2-dimensional plane which captures most of the volume of +TIP bodies to achieve a time resolution of around 31 frames per second. Our highly time-resolved acquisitions allow us to distinguish when nanoscale condensates first interact (Figure 1C) and we leverage this information to provide an estimate for their length scale (Figure 2). We find remarkably that +TIP bodies can reach 200-250nm diameter in size, which suggests that previous estimates of the number of Kar9 molecules based on fluorescence may be underestimates caused by fluorescent quenching^15^. The correspondence between our in vitro and in vivo measurements of material properties is striking (Figure 4B). Future work should attempt to dissect the contributions of viscosity and surface tension. Nonetheless, in vitro measurements of Bik1 condensates^30^, which are likely less dense and viscous than KBB droplets and +TIP bodies^15^, have viscosity which is much higher than estimates of the viscosity experienced by astral microtubules of the budding yeast cytoplasm^34^. This suggests that the simplifying assumptions made to calculate inverse capillary velocities are valid Figure 4A).

Microtubule depolymerization releases energy stored by polymerization and is therefore thought to be harnessed by cells to do work^2^. In the case of the budding yeast +TIP body this force is harnessed to move the spindle, which ensures correct inheritance of the chromosomes into mother and daughter cells. We propose that capillary forces are a parsimonious model to explain how budding yeast +TIP bodies track shrinking microtubules. In this model, a balance between cohesion, driven by multivalency, and adhesion, driven by affinity for microtubules, defines a preferred contact angle with the microtubule shaft. All the capillary velocities we measured in living cells in this study (range from 0.06µm/s -1.72µm/sec, Figure 4B) were faster than microtubule depolymerization velocities (highest velocity = 0.06µm/sec, Figure 5E). Therefore, the timescale on which tubulin heterodimers are lost from the polymer is slower than the timescale at which the +TIP body can relax to its preferred contact angle. We speculate that as protofilaments curve outward, the +TIP body moves down the shaft of the microtubule. Since KBB droplets are viscoelastic (Figure 4E-F), we propose that an elastic modulus of the +TIP body can transmit this force to the cortex through links made by Myo2 motors at its surface (Figure 3). We also speculate that capillarity may contribute to the ubiquitous tracking of growing microtubules by +TIP networks^2^.

Microtubule plus-end dynamics are intrinsically stochastic, yet groups of microtubules become synchronized to achieve critical cellular processes; for example, metazoan chromosome segregation and nuclear congression during budding yeast mating^2,35^. In this study, we observe metaphase +TIP bodies undergo fusion and fission and even undergo several consecutive fusion events (Figure 2D). This strongly suggests that some +TIP bodies harbor multiple synchronized plus-ends and that budding yeast spindle positioning also harnesses the force of several microtubules simultaneously. Since very little is known about how microtubules are synchronized, this is an exciting direction of further study which the budding yeast spindle positioning mechanism is well-suited to dissect. We speculate this ability may be a feature created by the +TIP body: a specialized nano-environment. In agreement with this, we found that +TIP bodies can reach a size 8-fold greater than a single microtubule diameter, around 200-250 nm (Figure 2), which could certainly accommodate several plus-ends. We were also intrigued to find that the microtubule depolymerization velocity is faster during cortical anchoring events in Kar9 oligomerization mutants (Figure 5E), which we speculate is a result of the less dense (Figure S3) local environment.

Our results are the strongest direct evidence that +TIP condensates are bona fide fluids with the potential to enact capillary forces in cells. Ou results suggest these properties can harness forces of dynamic microtubules, enabling them to work together. Our in vivo measurements have the potential to guide studies on other nanoscale condensates, including other +TIP condensates, which represent an important frontier in the field of biomolecular condensates.

## Acknowledgements

We thank the members of the Vogel and Barral labs, the participants of the Physical Basis of Cellular Memory and Adaptation workshop, and Eric Dufresne for helpful discussions. We thank Khalid Al-Naemi for constructing strains which use BDFP tag on Spc42 and Randal Halfmann for providing plasmids with BDFP for use in yeast. We thank the Steinmetz lab for reagents related to protein purification. K.M. is supported by a Maximilan Eivaskhani Memorial Fund from the Centre for Research in Biological Structure (CRBS).

This work is supported by grants from the Swiss National Science Foundation (31003A-105904 and Sinergia CRSII5_189940) and from SystemsX.ch (RTD grant #2012/192 TubeX) to Y.B. and grants from the CRBS (Blue Sky program), the Canadian Institutes of Health Research (PJT-1666078), the Natural Sciences and Engineering Research Council of Canada (RGPIN-2020-05187) and the Canadian Foundation of Innovation (JELF-40769) awarded to J.V.

## Author Contributions

Conceptualization, K.M., J.V., Y.B., and S.M.M.; Formal analysis, K.M., A.Z., and S.M.M.; Funding acquisition, J.V. and Y.B.; Investigation, K.M., S.M.M., M.C., A.Z., and M.W.; Methodology, K.M., S.M.M., J.V., and A.Z.; Resources, Khalid Al-Naemi, Randall Halfmann, Michel Steinmetz; Supervision, K.M., J.V., and Y.B.; Visualization, K.M., J.V., S.M.M., and M.C.; Writing – original draft, K.M.; Writing-review & editing, K.M., J.V., S.M.M., and Y.B..

## METHODS

### KEY RESOURCES TABLE

| Reagent or Resource | SOURCE | IDENTIFIER |
| --- | --- | --- |
| **Experimental Models: Organisms/ Strains** |  |  |
| Kar9-mNG; Spc42-mK2 | Meziane et al.<sup>21</sup> | YV2380 |
| Kar9-triple-mNG; Spc42-mK2 | This study | YV3212 |
| Kar9-mK2; Kar9-mNG | This study |  |
| Kar9-mK2; Myo2-998-GFP; Spc42-BDFP | This study |  |
| Myo2-998-GFP; CFP-tub1 |  |  |
| **Software and Algorithms** |  |  |
| Custom Matlab | This study |  |
| ImageJ | NIH | https://imagej.nih.gov/ij/ |
| TrackMate Version 7 |  |  |

### RESOURCE AVAILABILITY

#### Lead Contact

Further information and requests for resources and reagents should be directed to and will be fulfilled by the lead contact, Jackie Vogel.

#### Materials Availability

Further information and requests for resources and reagents should be directed to and will be fulfilled by the lead contact, Jackie Vogel.

### EXPERIMENTAL MODEL AND SUBJECT DETAILS

#### Yeast strain construction

All yeast are descendants of BY4741, BY4743, or S288C. All yeast strains in this study are found in key resources table.

### METHOD DETAILS

#### Plasmid preparation, recombinant protein expression and purification for in vitro time-resolved microscopy

In vitro time-resolved microscopy data of protein condensates from Meier et al 2023 was reanalyzed to determine fusion times and condensate viscoelasticity and aging in this study. In short, to produce the recombinant proteins used in this assay, full-length hexa-histidine tagged wild-type or self-oligomerization interface mutant (Phe292Ala, Leu347Ala, Arg237Ala, Asn241Ala, Tyr366Ala, Arg367Ala) Kar9 (UniProt ID P32526) were cloned into ACEMBL vector pACE, full-length hexa-histidine tagged Bik1 (UniProt ID P11709) into PSTCm1, as in Olieric et al.^36^, and full-length Bim1 (UniProt ID P40013) into pET3d (Invitrogen)^37^. These plasmids were transformed into BL21(DE3) *E. coli* cells for expression, which were subsequently grown in liquid cultures of LB containing appropriate antibiotics while shaking at 37°C until optical density at 600 nm of 0.6 was reached. Protein expression was induced by addition of 0.75 mM isopropyl-β-D-thiogalactoside at 20°C overnight. Cells were lysed using the sonication method or high-pressure homogenizers (Avestin Emulsiflex-C3 High Pressure Homogenizer) in 20 mM Tris–HCl (pH 7.5), supplemented with 500 mM NaCl. For co-purification of Kar9 and Bim1, cells expressing each individually were mixed at equal ratio prior to lysis. Cell debris was removed by centrifugation and lysates were further cleared by filtration through 0.45 μm filters before purification of hexa-histidine tagged proteins (Bik1) or protein complexes (Kar9-Bim1) by a]inity chromatography on Ni2^+^-Sepharose columns (Cytiva) at 4°C according to the manufacturer’s instructions. Proteins were further purified by size exclusion chromatography using Superdex-75 or −200 (Cytiva) at 4°C and concentrated by Vivaspin (Sartorius) concentrators at 4°C (Kar9-Bim1) or 25°C (Bik1). The purity of recombinant proteins was confirmed by Coomassie-stained SDS–PAGE, the identities of the proteins were assessed by mass spectral analyses, and concentrations were assessed by absorption at 280 nm.

For time-resolved microscopy, Bik1 and wild-type or mutant Kar9-Bim1 complex were mixed at 4X concentration and 500 mM NaCl before dilution to 10 μM and 220 mM NaCl to induce condensation in a Nunc Lab-Tek II chambered eight-well coverglass in a total volume of 30 μl. Microscopy was started within 30s after dilution. To verify enrichment of His-tagged protein in droplets, Atto-488-NTA was used at 1 μg ml^−1^ concentration.

Samples were imaged on a DeltaVision Personal microscope (Cytiva) at 60× magnification using transmission and FITC channels in a single plane close to the coverglass surface.

#### Extraction of yeast lysate

Yeast cells harboring Myo2-998-GFP and CFP-Tub1 were grown overnight at 25°C while shaking. Cells were harvested at log phase by spinning down at 700g for 5 minutes. Cell pellet was washed with 1 mL cold PBS by spinning at 21300 rcf for 1 minute at 4°C in a screw cap tube. To a cell pellet made from 20 ml of culture volume having OD_600_ of 0.5, 200 µL of glass beads measured with an Eppendorf (0.5 mm Glass Beads, Catalog# 11079105, BioSpec Products) were added along with 400 µL of cold lysis buffer (50mM Tris, 500mM NaCl, 0.5 mM EDTA, 1 mM MgCl_2_, 0.2% Triton X-100, pH 7.5) freshly supplemented with protease cocktail inhibitor (cOmplete EDTA-free protease inhibitor cocktail tablets, Roche Diagonistics). Cells were opened at 4°C in a bead beater (MP FASTPREP-24^TM^ 5G) at 6m/s, 2 cycles each of 50 sec with 2 minutes break in between. Lysate was cleared at 21300 g for 15 minutes at 4°C. Supernatant was separated and directly used for microscopy experiments.

#### Droplet partitioning assay

Kar9, Bim1 and Bik1 purified with 500 mM NaCl were premixed with yeast lysate, with purified protein concentration of 10 µM and yeast lysate comprising 75% of the intended premix volume. Phase separation was induced by diluting the NaCl concentration to 250 mM NaCl using buffer containing 20 mM Tris in a eight-well chambered Coverglass (Lab-Tek II Chambered coverglass, Catalog# 155409). Droplets were imaged on a Delta Vision microscope (Cytiva) at 100× magnification using transmission and FITC channels in a plane close to the coverglass surface.

#### Live-cell structured illumination microscopy

Yeast cells were grown overnight in synthetic complete (SC) media at 25C with shaking to saturation and the morning of imaging diluted to OD_600 0.2 SC supplemented with 2mM ascorbate, then cultured for ∼2 hours to achieve mid-log phase growth (∼OD_600 0.4). All imaging was performed at 25C. Cells were prepared for imaging as described previously^31^. In brief, cells are pelleted by spinning at 500 rpm for 30 seconds and washed once in SC supplemented with 2mM ascorbate. Cells are then spun again at 500 rpm for 30 seconds and supernatant is mostly removed to concentrate cells. 4 ul of resuspended concentrated cells are pipetted onto a clean slide and a coverslip (Carl Zeiss Cover Glass, High Performance; 0.170 mm +/-0.005 mm) is applied. Each slide is used for a maximum of 20 minutes. All imaging is done using the Elyra 7.2 (Carl Zeiss) equipped with an AxioObserver 7 with motorized focus drive, Optovar 1.6x, Scanning stage Piezo, Plan-Apochromat 63x/1.40 Oil DIC M27 objective (Carl Zeiss), and pco.edge sCMOS (Version 4.2 CL HS) cameras, with DuoLink adapter to connect cameras for simultaneous dual-color imaging. Channel alignment was performed using Zen Black 3.0 and multicolor calibration slide (Carl Zeiss).

To visualize +TIP bodies and spindle position over time, 3-phase linear structured illumination (Zeiss apotome mode) time series were acquired using laser 488nm/500mW and 561nm/500mW laser lines both at 7%. Red and green channels were captured simultaneously using LBF 405/488/561/642 using a dichroic with emission filters BP 570-620+LP 655 and BP 420-480+BP 495-550. Stacks were acquired with 0.3 um space between slices, with LEAP mode processing final voxel size of 0.1, 0.031, 0.031 µm for a final stack depth of 3.9 um. Acquisition of each stack takes ∼1.6 seconds, and 20 timepoints were acquired with a 3.82 second timestep. Cells appearing to be in late metaphase or anaphase were identified based on cellular morphology and cropped using the Fiji roi manager or Zen Blue.

To capture fusion events in live cells, 2D time series were acquired for ∼20 seconds using lattice structured illumination with 9 phases and 25 ms exposure time per phase.

Cells were illuminated with 488nm/500mW and 561nm/500mW laser lines at 5.0% and 2.0% respectively. Both red and green channels were captured simultaneously using beam splitter LBF 405/488/561/642 and a dichroic with emission filters BP 570-620+LP 655 and BP 420-480+BP 495-550. Zen Black 3.0 software (Zeiss) was used to reconstruct super-resolution images with peak intensity scaled to raw phase images. Burst processing (Zeiss) was used to achieve an effective frame rate of 31 frames per second. Candidate fusion events in which two foci appear to merge were cropped manually using the Fiji roi manager.

To capture +TIP body composition, near-simultaneous 3-color imaging was achieved using frame fast mode for rapid switching between laser lines at each slice of the stack using 25 ms exposure per laser line. 60% 561nm, 40% 642 nm, and 30% 488 nm illumination was used for diploid cells (Kar9-mK2, Kar9-mNG), 40% 561nm, 40% 642 nm, and 50% 488 nm illumination was used for haploid cells (Kar9-mK2, Myo2-GFP, Spc42-BDFP). Near-simultaneous imaging of three channels was acquired using the beam splitter LBF 405/488/561/642and dichroic with emission filters BP 570-620+LP655 followed by BP 420-480+BP 495-550 and BP 570-620+LP655 simultaneous acquisition. These acquisitions used 3 phase apotome mode z:15 with total stack 2.8 um, and acquisition of each stack takes ∼1.6 seconds. For super-resolution reconstructions, 0.2 um sections were processed to be .067 nm .031, .031 voxel size using LEAP mode reconstruction (Zeiss).

### QUANTIFICATION AND STATISTICAL ANALYSIS

#### In vitro droplet fusion analysis

Fusion events in which two droplets merge were cropped manually using Fiji roi manager. The ‘moment of fusion onset’ was taken as the first frame of unambiguous uniaxial deformation after two droplets had come into contact. The shape in each frame from the ‘moment of fusion onset’ until the droplet had rounded up within error (by eye) was measured using the multi-point tool in Fiji. The Ellipse Long Axis (L) and Ellipse Short Axis (W) were saved. As in ^30^, these were used to measure the elliptical shape parameter A = (L-W)/(L+W) over time, which exhibits an exponential decay to 0 as the droplet rounds. The characteristic time tau of this decay was extracted by fitting A to the function a*exp(-(x/t))+b using the Matlab function **fit.** The final radius is taken as the final (L+W)/4 to determine the τ /R. For time-resolved τ/R analysis, the initial timepoint of fusion normalized to the start of the acquisition was used.

#### In vivo fusion analysis

Candidate fusion events were cropped in xy to a window of a few microns and in time to a section of 100-300 frames (∼3-10 seconds) in Fiji using the roi manager. The Fiji plugin Trackmate using the trainable Weka detector was then used to segment +TIP bodies by pixel classification and trajectories were established using the Trackmate LAP tracker. Tracks for objects within the same roi but not of interest were manually removed. Object parameters of Maximum intensity, Sum intensity, X position, position, Ellipse Long Axis (L), Ellipse Short Axis (W), and Ellipse Aspect Ratio were saved. Tracks with sum intensities which change by more than 2.5 fold, indicating +TIP bodies are moving in or out of focus, were not analyzed (excluded: n=5 WT events, 5 cells, n = 5 Kar9-αβγ^-^ events, 5 cells). L and W were used to measure the elliptical shape parameter A = (L-W)/(L+W) over time using Matlab. The majority of fusion events achieved an ellipticity of 0.1 (n= 13/14 WT events, n=14/17 Kar9-αβγ^-^ events). Based on sum intensity and shape parameter trajectories, trajectories were further cropped in time to span the initial detection of one anisotropic shape until Ellipticity of <0.1 is achieved. Within this trajectory, the ‘moment of fusion onset’ was determined manually by looking for the changepoint in the Ellipticity, Long Axis (L), and Ellipse Aspect Ratio (L/W) trajectories, moving backward in time from the achievement of Ellipticity of <0.1. As with the in vitro analysis, the characteristic time tau was extracted by fitting A from the ‘moment of fusion onset’ to the end of the trajectory using the function a*exp(-(x/t))+b. The final radius is taken as the mean (L+W)/4 in the final 10 frames (equalling about 0.3 seconds), and this was used as the R in the calculation of τ/R.

#### Distance at fusion onset analysis

At the timepoint previously determined as the moment of fusion onset, a 1 µm linescan of 3 pixel width was taken across the axis of fusion. The intensity values along this line were saved and analyzed using a custom Matlab script which employs the fit function with the fittype ‘gauss2’ to fit the intensity value curve to a sum of two gaussians. The distance between the means was reported as the peak distance at fusion onset.

#### Radial intensity profile analysis

Using the Fiji plugin Trackmate with Laplacian of Gaussian detector was used to determine the centroid pixel in red channels in 3D with a lower bound for quality of points set by the threshold 28. Invalid localizations (detections outside of cells or at the edges of the FOV) were removed manually or by increasing the quality threshold for individual images. A custom Fiji macro (adapted from [citation]) was used to measure the normalized radial intensity profile from 0 to 0.5 microns. Localizations <0.5 micron from any edge of the FOV were excluded from this analysis. Each profile was binarily classified by whether the maximum intensity along the normalized radial profile occurred at the center or outside of the center.

#### Fast anaphase analysis

Using the Fiji plugin Trackmate with Laplacian of Gaussian detector and LAP tracker, tracks for the old and new spindle pole bodies were determined. Spindle length was measured as the 3D distance between old and new spindle pole bodies. The spindle growth rate was taken by the slope of the line of best fit through the spindle length values over time. Fast anaphase cells were identified if they met either criteria: 1) Having an initial spindle length >1.8 um and growth rate > 0.2 micron/minute or 2) Having the spindle pole body closest to the bud neck at the initial frame pass through the bud neck during the acquisition.

#### Spindle pole body regression analysis

Progression of the spindle pole into the bud was measured in Fiji by using the line tool to draw a line parallel to the angle of the bud neck and intersecting the pole at the initial frame. If during the acquisition, the pole crosses this line and becomes closer to the bud neck, this cell is counted as having a regression. Cells with poles that maintain or advance their position into the bud are considered progressed.

#### Depolymerization pulling event analysis

Candidate pulling events during which the bud adjacent or residing spindle pole body moves towards a cortical focus of Kar9 were identified using the 3D viewer plugin and 3D Sum projections in Fiji. For each event, Kar9 trajectories were tracked using the Fiji plugin Trackmate. The 3D distance to spindle pole bodies was used to confirm depolymerization during events. Kar9 trajectories were limited to moving less than 0.2 µm between each frame. Events lasting at least 3 frames (8s; 3.8s between frames) were segmented. For each trajectory, duration, speed of depolymerization, and type of event end was quantified.

---

**Figure S1.**
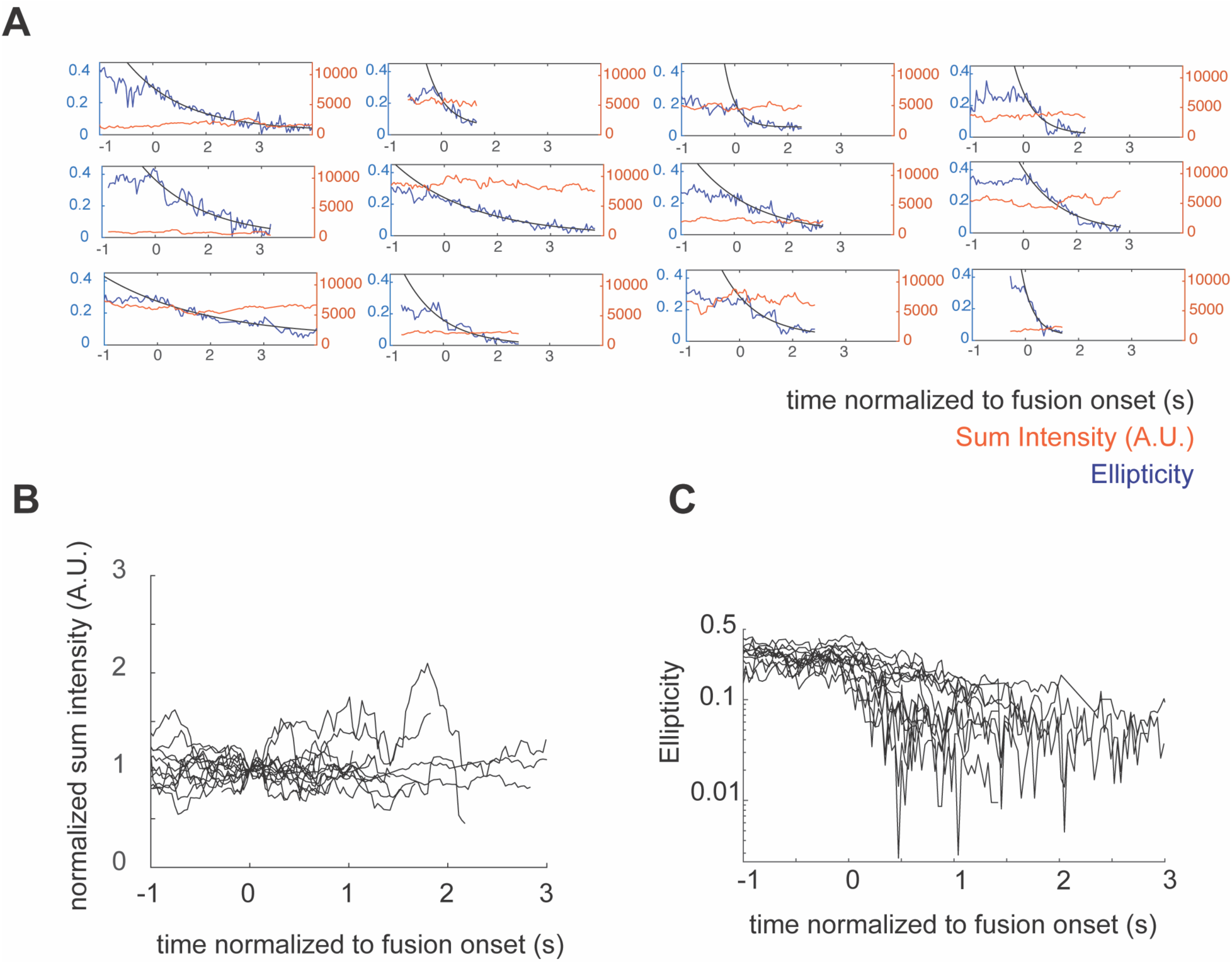
Raw trajectories of +TIP body shape recovery dynamics in vivo, related to Figure 1. (A) Plots of ellipticity and sum fluorescent intensity (A.U.) over time for individual wild type cells. Ellipticity is defined as (L-W)/(L+W) where L = length and W = width. (B) Plot of normalized sum fluorescent intensity (A.U.) over time for wild type cells. (C) Plot of ellipticity over time for wild type cells.

**Figure S2.**
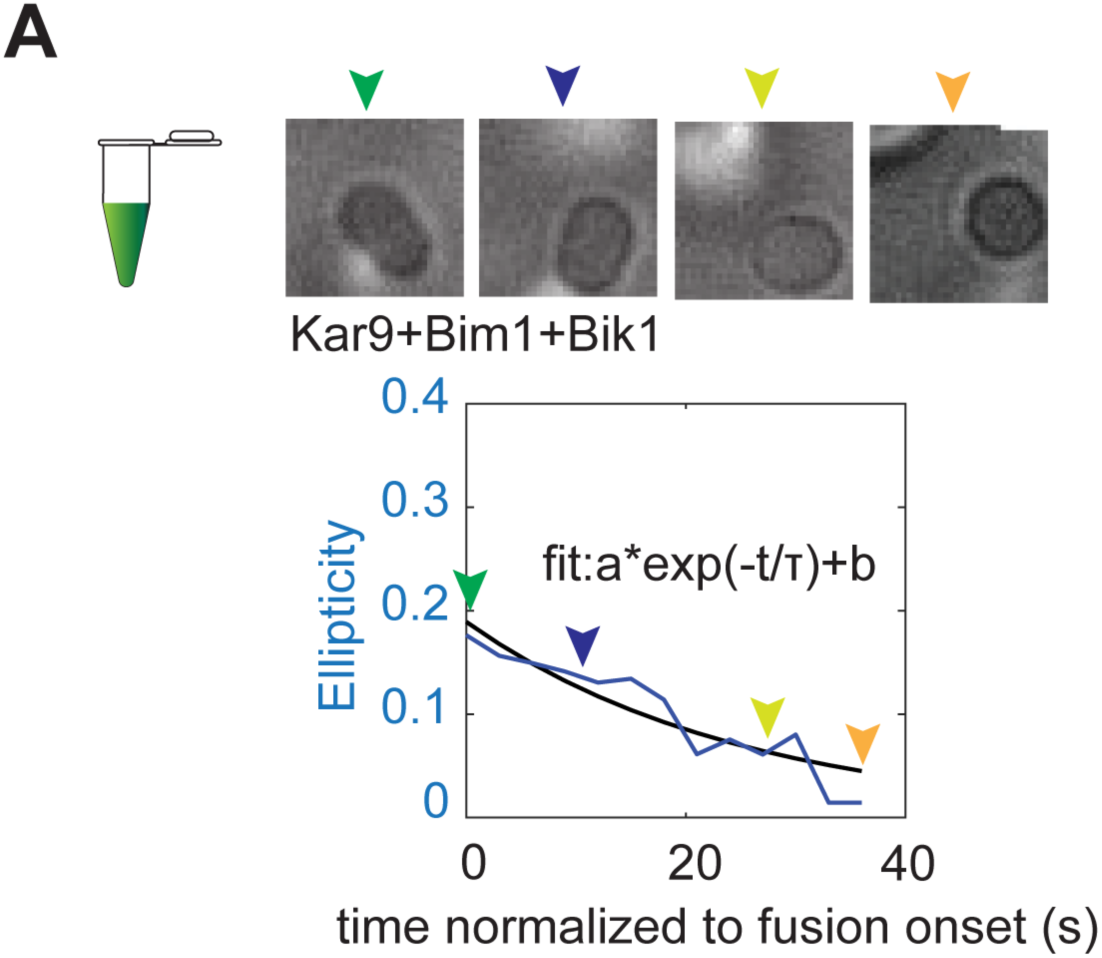
In vitro fusion analysis, related to Figure 4. (A) KBB droplet merging event. Scale bar 1 µm. Plot of Ellipticity (as in Figure 1, Figure S1) over time and exponential fit to extract the characteristic time t.

**Figure S3.**
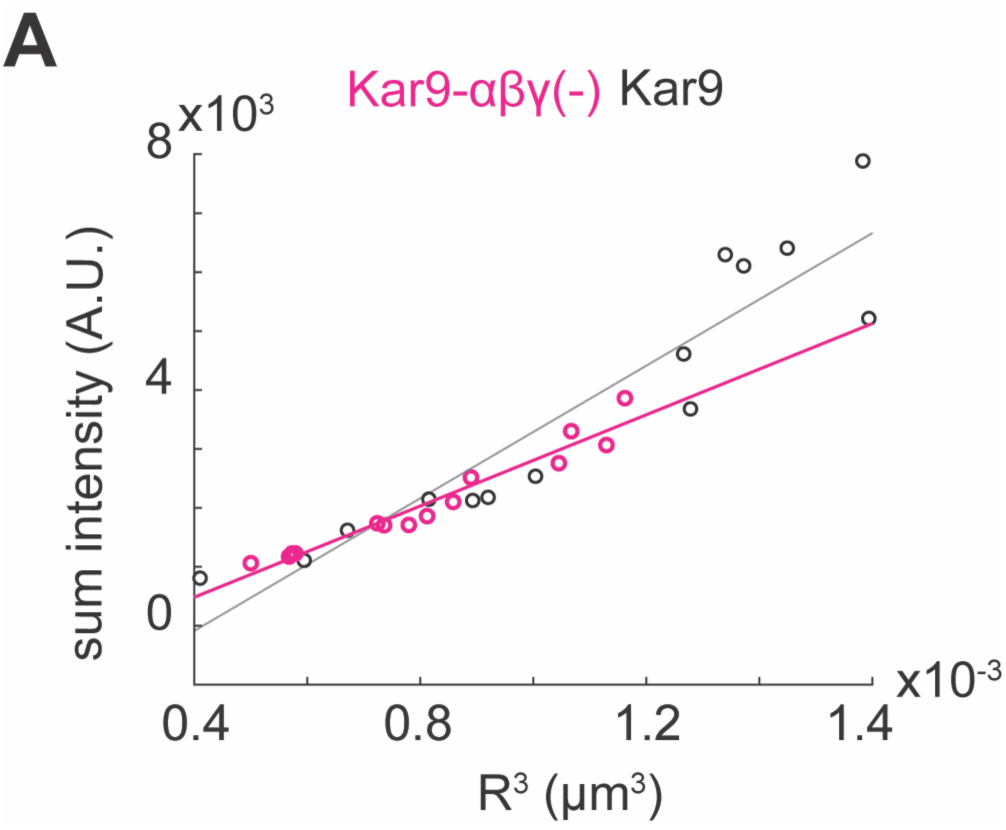
+TIP body density in vivo, related to Figure 4. (A) Sum intensity vs R^3^ for +TIP bodies resulting from merging events in wild type cells and Kar9-αβγ(-) cells.

**Figure S4.**
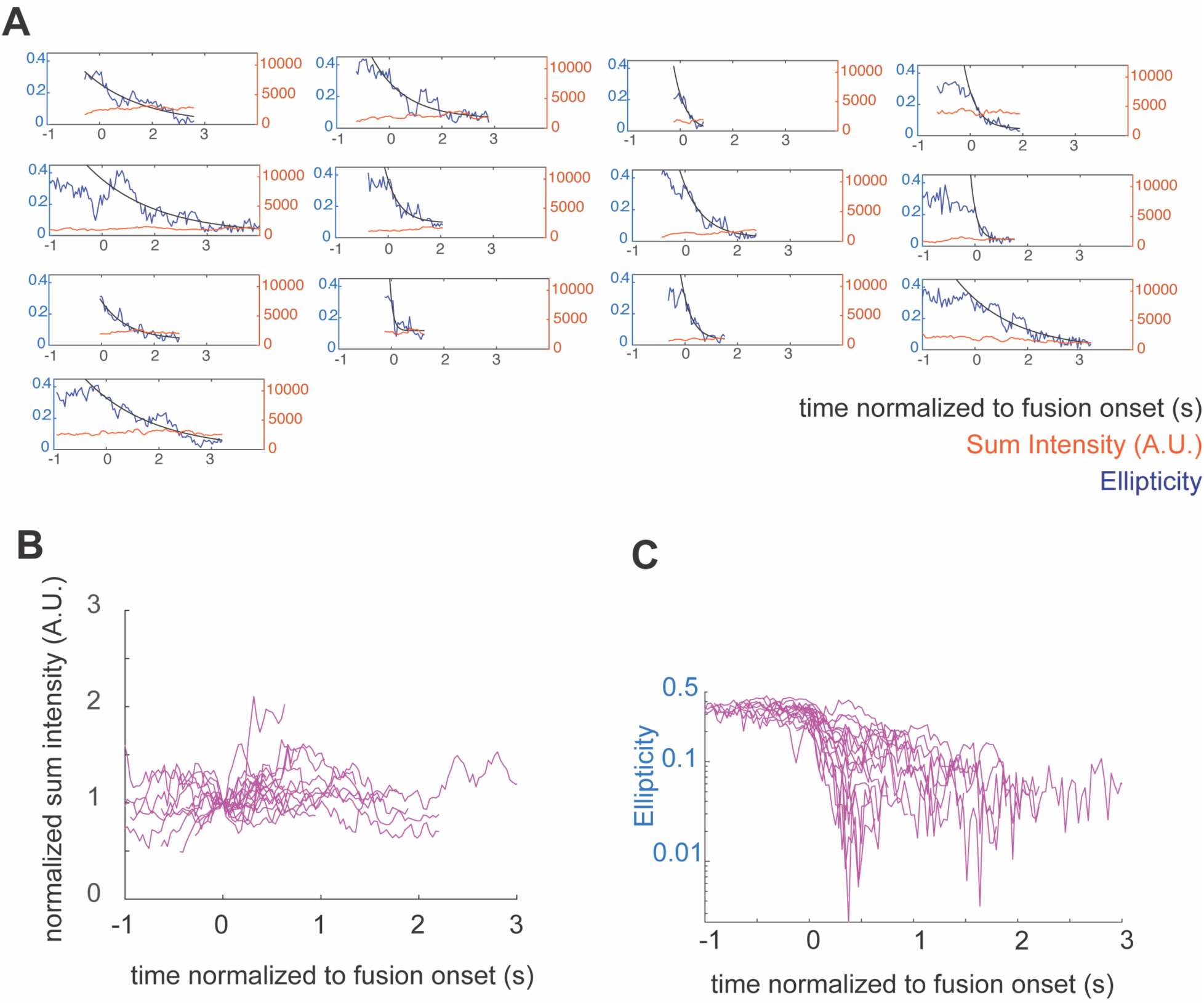
Raw trajectories of mutant +TIP body shape recovery dynamics in vivo, related to. **Figure 4** (A) Plots of ellipticity and sum fluorescent intensity (A.U.) over time for individual Kar9-αβγ(-) cells. Ellipticity is defined as (L-W)/(L+W) where L = length and W = width. (B) Plot of normalized sum fluorescent intensity (A.U.) over time for Kar9-αβγ(-) cells. (C) Plot of ellipticity over time for Kar9-αβγ(-) cells.

**Figure S5.**
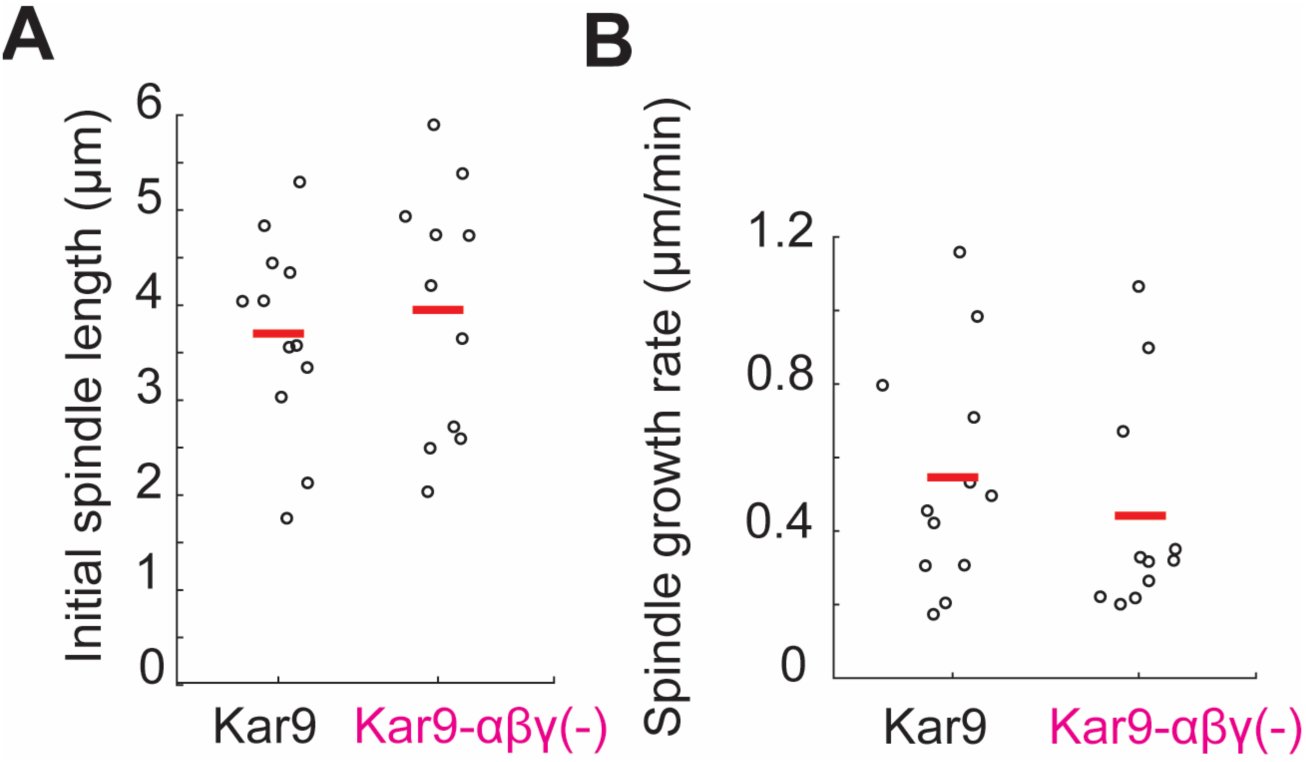
Initial spindle length and spindle growth rate, related to Figure 5. (A) Initial spindle length. Measured by 3D distance between spindle pole bodies. (B) Spindle growth rate. Measured using a linear fit of the trajectory of the 3D distance between spindle pole bodies over the acquisition.

